# Early-life gut dysbiosis linked to mass mortality in ostriches

**DOI:** 10.1101/841742

**Authors:** Elin Videvall, Se Jin Song, Hanna M. Bensch, Maria Strandh, Anel Engelbrecht, Naomi Serfontein, Olof Hellgren, Adriaan Olivier, Schalk Cloete, Rob Knight, Charlie K. Cornwallis

**Author notes:** **Corresponding author**, Elin Videvall.

## Abstract

Dysbiosis in the vertebrate gut microbiome has been associated with several diseases. However, it is unclear whether particular gut regions or specific time periods during ontogeny are responsible for the development of dysbiosis, especially in non-model organisms. Here we examine the microbiome associated with dysbiosis in different parts of the gastrointestinal tract (ileum, caecum, colon) in a long-lived bird with high juvenile mortality, the ostrich. Individuals that died of gut disease (n=68) had substantially different microbial composition from age-matched controls (n=50) throughout the gut. Several taxa were associated with mortality (Enterobacteriaceae, Peptostreptococcaceae, Porphyromonadaceae, *Clostridium*) and some with survival (Lachnospiraceae, Ruminococcaceae, Erysipelotrichaceae, *Turicibacter*). Repeated faecal sampling showed that pathobionts were already present shortly after hatching and proliferated in individuals with low diversity, resulting in mortality weeks later. The factors influencing seeding of the gut microbiota may therefore be key to understanding dysbiosis and host development.

## Introduction

The composition of the microbial community in the gastrointestinal tract of animals (‘the gut microbiome’) is extremely important for host fitness and health (McFall-Ngai *et al.* 2013). Imbalances in the gut microbiome, commonly referred to as gut dysbiosis, has been widely associated with a variety of gut and autoimmune diseases such as type 1 diabetes, Crohn’s disease, inflammatory bowel disease, ulcerative colitis, and multiple sclerosis (Wu & Wu 2012; Gevers *et al.* 2014; Machiels *et al.* 2014; Berer *et al.* 2017; Duvallet *et al.* 2017). Dysbiosis is typically characterized by loss of beneficial microorganisms, proliferation of pathobionts (opportunistic microorganisms), and loss of overall microbial diversity (Sekirov *et al.* 2010; Petersen & Round 2014). Transplants of gut microbiota from mice with gut disease have been shown to result in similar disease symptoms in recipients, suggesting a strong causal effect of gut dysbiosis on host health (Garrett *et al.* 2010; Vijay-Kumar *et al.* 2010). Inflammation of the gastrointestinal tract is often associated with gut dysbiosis, which in turn alters the intestinal mucus layer and epithelial permeability resulting in increased susceptibility to infection, sepsis, and organ failure (Zimmermann *et al.* 2012; Latorre *et al.* 2015; Klingensmith & Coopersmith 2016).

When and where imbalances in gut microbiota originate is unclear. The diversity and composition of microbes differ markedly across different regions of gut (Zhang *et al.* 2014; Donaldson *et al.* 2015) and it is possible that certain regions may act as sources of pathobionts, radiating out to disrupt other parts of the gut. For example, certain regions might be more susceptible to pathogenic overgrowth due to low microbial diversity and reduced resilience (Sommer *et al.* 2017). Alternatively, dysbiosis may occur throughout the gastrointestinal tract, or develop from diverse communities that harbour more pathobionts. Pin-pointing when groups of bacteria start to proliferate in different regions of the gut has been difficult because most studies have used cross-sectional sampling (one sample per individual). As a result, it remains unclear whether bacteria associated with dysbiosis are always present in low abundance, or whether dysbiosis is linked with a sudden influx of foreign microbes from an external source.

An additional problem has been to establish whether certain groups of bacteria are consistently involved in dysbiosis across diverse host species. The vast majority of microbiome studies, and specifically those on dysbiosis, have focused on humans and laboratory mice (Petersen & Round 2014). This research has shown that certain bacterial taxa seem to be routinely associated with dysbiosis across species and individuals. For example, in inflammatory bowel disease, one of the most common indicators of dysbiosis is elevated levels of Enterobacteriaceae (Gammaproteobacteria) (Lupp *et al.* 2007; Garrett *et al.* 2010; Hughes *et al.* 2017), and a reduction of Ruminococcaceae and Lachnospiraceae (Clostridia) (Antharam *et al.* 2013; Duvallet *et al.* 2017). Whether these patterns extend across more distantly related species and outside laboratory settings is unclear, especially for non-mammalian organisms.

In this study we utilise the ostrich (*Struthio camelus*) as a new vertebrate host system to understand patterns of gut dysbiosis and its effect on mortality. Ostriches suffer from exceptionally high and variable mortality rates at rearing facilities during their first three months of age (Verwoerd *et al.* 1999; Cloete *et al.* 2001). While the causes of mortality are mostly unknown, several candidate pathogens associated with enterocolitis have been reported, for example *Escherichia coli*, *Campylobacter jejuni*, *Pseudomonas aeruginosa*, *Salmonella* spp., *Klebsiella* spp., and multiple *Clostridium* spp. (Frazier *et al.* 1993; Huchzermeyer 1999; Shanawany & Dingle 1999; Verwoerd 2000; Keokilwe *et al.* 2015). However, whether variation in mortality is due to infection of specific pathogens or the result of microbiome dysbiosis has not yet been established. The studies investigating causes of mortality in ostrich chicks have so far utilized methods involving bacterial culture or species-specific DNA primers. These methods can be useful to detect the presence of targeted microorganisms, but searching for a particular culprit may yield ambiguous answers if pathobionts exist in the normal gut microbiota of the host and only exhibit pathogenic tendencies when the community is disturbed (Chow *et al.* 2011). In addition to a high mortality rate, ostriches exhibit large variation in microbial composition between individuals and across gut regions (Videvall *et al.* 2018). Because these animals have only been reared in captivity for a very short time relative to other farmed animals (Cloete *et al.* 2012), they exhibit several of the advantages of a wild study system (high genetic variation, non-domesticated social groups, reared under semi-natural conditions) while still allowing for standardised conditions and ease of sampling.

Here we evaluate dysbiosis-associated mortality in 68 individuals that died from suspected gut disease within three months after hatching (referred to as “diseased”) and compare it to 50 individuals that were euthanized as age-matched healthy controls (referred to as “controls”). The microbial composition of the ileum, caecum, and colon were characterized to determine the pattern of dysbiosis in different regions of the gastrointestinal tract. Faecal samples collected at 1, 2, 4, and 6 weeks of age from the same 118 individuals plus four additional individuals that survived the whole period (n = 122) were analysed to investigate when dysbiosis-related features emerge. Finally, samples from food, water, and soil were examined to evaluate potential sources of pathogenic bacteria. We use these results to discuss whether certain characteristics of gut dysbiosis may be present across a diverse range of host species.

## Results and discussion

### Mortality and dysbiosis in different gut regions across ontogeny

Mortality of juvenile ostriches occurred throughout the entire 12-week period but was highest between four and eight weeks of age, with a peak at six weeks of age (Figure 1b). Individuals with disease followed the growth curve of all other individuals, but rapidly dropped in weight prior to death (Figure 1c-d). The cause of the weight reduction is unknown, but diseased individuals were observed to stop eating and drinking, and in some cases suffered from diarrhoea, so dehydration and wasting are likely explanations. In total, 40% of all chicks died (excluding controls) and post-mortems of diseased and control individuals revealed that mortality was associated with extensive inflammation of the gastrointestinal tract (Figure 1e; Figure S1). The gut inflammation scores of diseased individuals (mean ± SD for ileum = 4.1 ± 1.0, caecum = 3.0 ± 1.3, colon = 3.0 ± 1.2) were substantially higher than those of control individuals (ileum = 1.4 ± 1.0, caecum = 1.0 ± 0.29, colon = 1.1 ± 0.45) (Figure S1).

**Figure 1.**
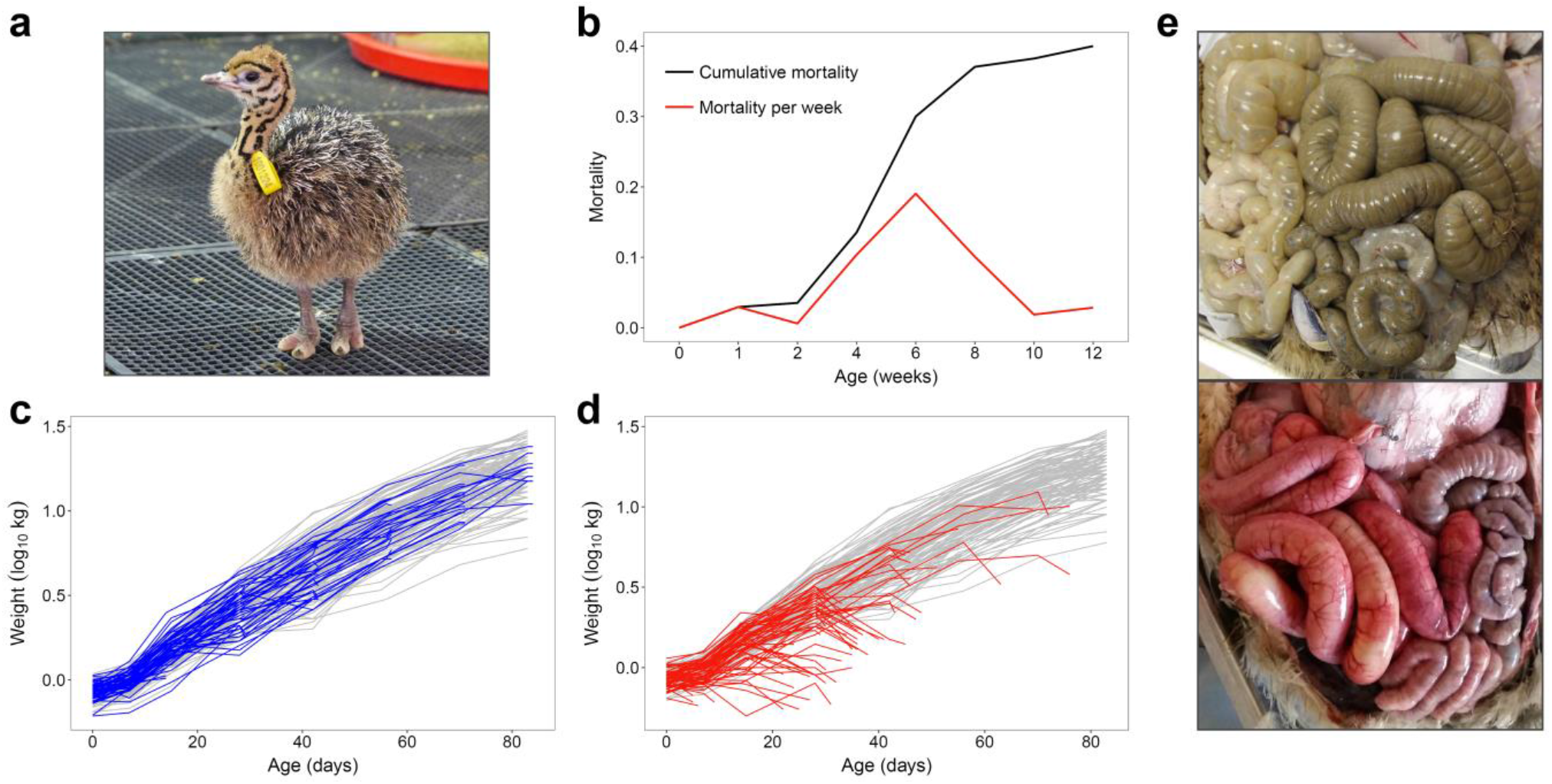
Mortality patterns of ostriches up to 12 weeks of age. (**a**) One of the ostrich chicks included in the study, here one week old. (**b**) The cumulative mortality and mortality rate per week. (**c, d**) Log-transformed weights over time of control individuals that were randomly selected for euthanization at weeks 2, 4, 6, 8, 10, and 12 (blue lines in **c**), and individuals that died of suspected disease (red lines in **d**). Grey lines illustrate weights of all other individuals that survived the whole period. (**e**) Photographs during dissection illustrating widespread gut inflammation in a diseased individual (bottom) compared to a control individual (top).

The structure of the microbiota was extremely different between diseased and control individuals in all three gut regions (Figure 2, Figure S2, Table 1). Specifically, we found significant differences in the microbial community distances between diseased and control individuals using both Bray-Curtis (BC) and weighted UniFrac (wUF) measures, controlling for age, sex, group, and time since death (Table 1). However, the magnitude of the differences between diseased and control individuals, measured with Bray-Curtis, decreased from the small intestine to the lower gut (ileum-caecum-colon); whereas the weighted UniFrac measures showed the opposite pattern, increasing in strength from the ileum to the colon (Table 1). The diseased individuals were more similar to each other in the ileal microbiome than the controls were to each other when using Bray-Curtis, but not weighted UniFrac distances (BC Multivariate homogeneity test of group dispersion (betadisper): F_1, 99_ = 13.9, p = 0.0003. wUF betadisper: F_1, 99_ = 0.6, p = 0.46) (Figures S3–S4). In contrast, the opposite was true for the caecum and colon (BC caecum betadisper: F_1, 105_ = 0.08, p = 0.79. BC colon betadisper: F_1, 106_ = 1.3, p = 0.25. wUF caecum betadisper: F_1, 105_ = 11.2, p = 0.001. wUF colon betadisper: F_1, 106_ = 11.4, p = 0.001) (Figure S3). These results indicate that the bacterial composition differed the most between diseased and control individuals in the ileum, but that the colon displayed the most phylogenetically distinct groups. Sex, group, and time since death had no significant effects on the microbiome in any of the gut regions using either of the distance measures (Table 1).

**Table 1.** PERMANOVA of microbiome dissimilarities across three gut regions.

|  | Ileum |  | Caecum |  | Colon |  |
| --- | --- | --- | --- | --- | --- | --- |
|  | BC | wUF | BC | wUF | BC | wUF |
| Disease | 15.5 *** | 8.2 *** | 10.8 *** | 11.7 *** | 7.9 *** | 18.2 *** |
| Age | 2.1 * | 0.8 | 5.7 *** | 3.0 ** | 7.1 *** | 5.8 *** |
| Age<sup>2</sup> | 1.6 * | 0.5 | 2.8 *** | 4.8 *** | 3.5 *** | 3.1 ** |
| Group | 3.1 | 4.9 | 2.8 | 1.9 | 3.0 | 2.2 |
| Sex | 0.6 | 0.3 | 1.1 | 0.8 | 1.1 | 0.8 |
| Time since death | 0.8 | 0.3 | 0.6 | 0.5 | 0.6 | 0.9 |

Effect sizes are displayed as $R^2$ values in percentage with the number of stars indicating level of statistical significance, \*\*\* $p < 0.001$ , \*\* $p < 0.01$ , \* $p < 0.05$ . BC = Bray-Curtis distances, wUF = weighted UniFrac distances.

**Figure 2.**
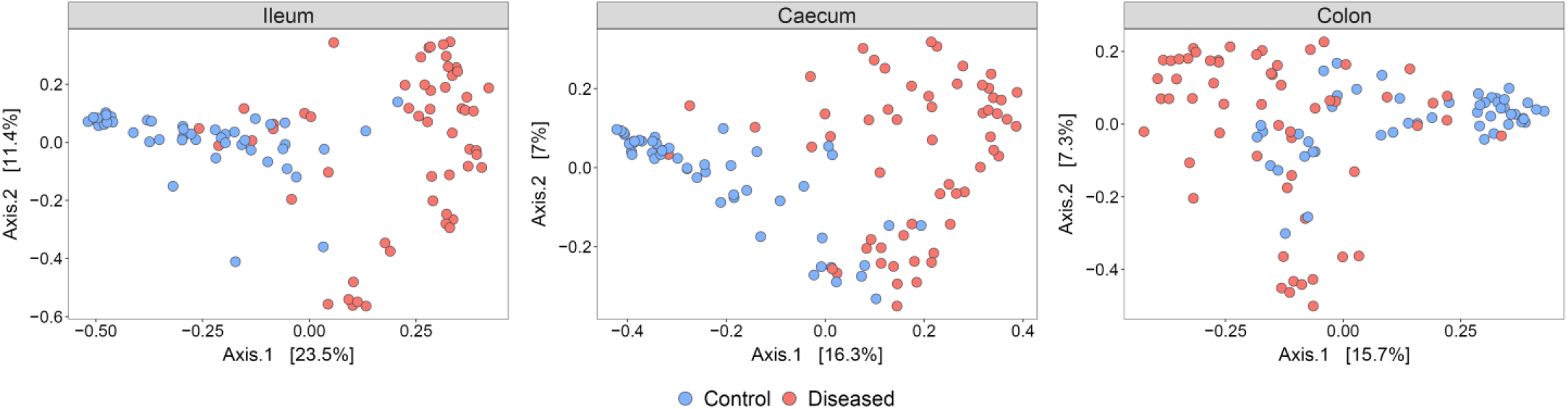
Principal Coordinates Analysis (PCoA) plots of Bray-Curtis dissimilarities between the microbiomes of control individuals (blue) and diseased individuals (red).

### Alpha diversity and age-specific dysbiosis

The microbial alpha diversity of diseased individuals was greatly reduced in all three gut regions in comparison to controls (GLM: ileum F_1, 99_ = 56.7, p = 2.5e−11; caecum F_1, 105_ = 16.1, p = 0.0001; colon F_1, 106_ = 61.5, p = 3.9e−12), controlling for age (Figure 3). These differences became less pronounced with age in the caecum and colon, but were consistent in the ileum regardless of age (GLM: ileum F_1, 97_ = 0.0001, p = 0.99; caecum F_1, 103_ = 10.2, p = 0.002; colon F_1, 104_ = 9.1, p = 0.003) (Table 1; Figure 3). Age had little effect on the development of the microbiome in the ileum even in healthy individuals (GLM: F_1, 98_ = 1.4, p = 0.23), whereas further down in the gut, diversity increased with age (caecum F_1, 104_ = 32.7, p = 1.0e−07: colon F_1, 105_ = 39.7, p = 7.2e−09; Figure 3). These results demonstrate that the microbial differences in diseased individuals are persistent across ages in the small intestine, but in the lower gastrointestinal tract these differences gradually diminish as the microbiome matures (see also Videvall *et al.* 2019).

**Figure 3.**
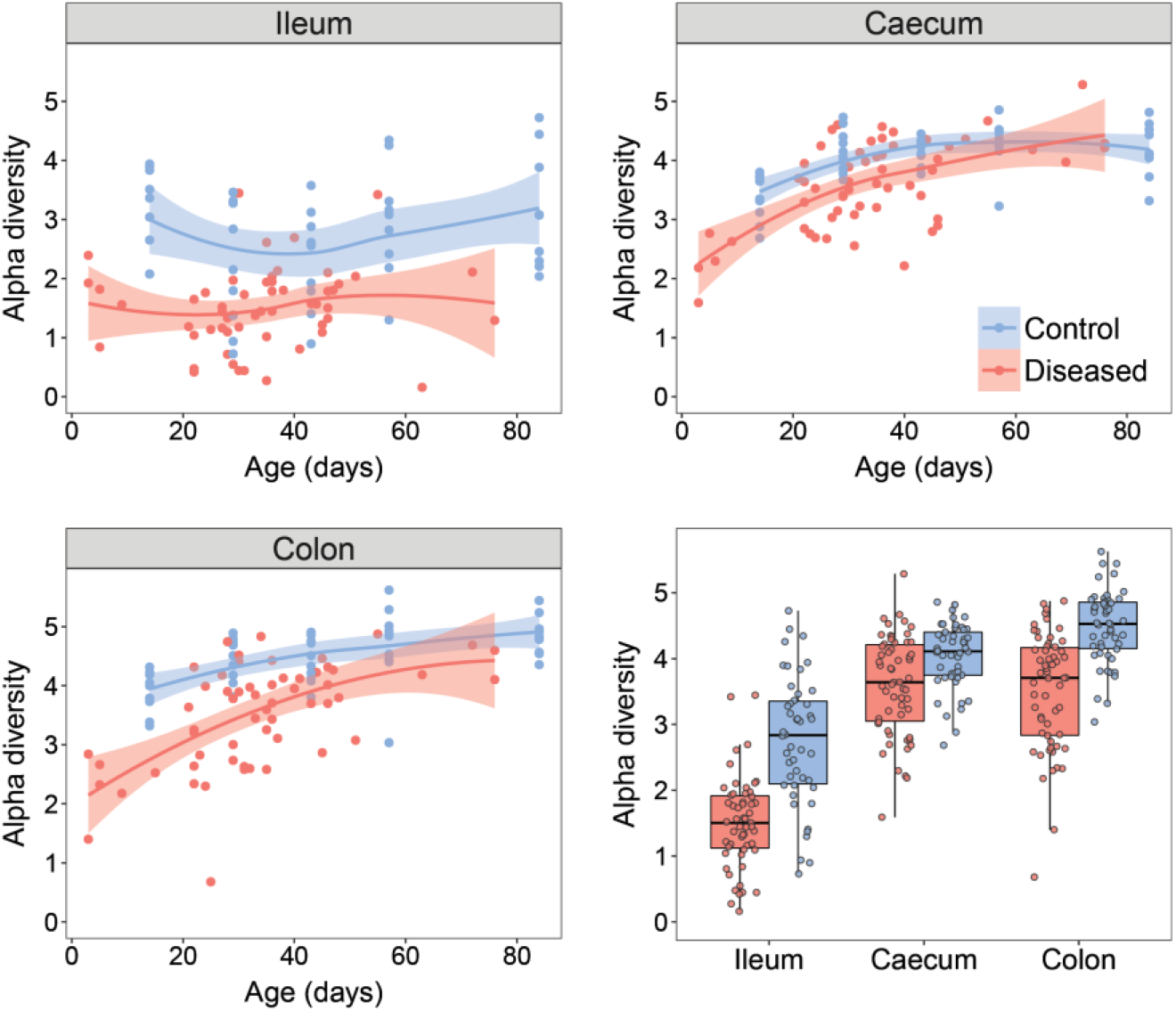
Alpha diversity (Shannon index) during development in the ileum, caecum, and colon. Control individuals are shown in blue and diseased individuals in red. Lines display the fitted local regression smoothing curves and shaded areas the 95% confidence interval. Bottom right panel shows all alpha diversity values together.

### Taxa associated with disease in the ileum

To better understand the microbial dissimilarities seen in diseased individuals, we evaluated the taxonomic composition of all gastrointestinal regions. The ileum showed the most striking evidence of dysbiosis (Figure 4). Control individuals had a diverse community of different bacterial classes in the ileum, whereas diseased individuals displayed a bloom of Gammaproteobacteria and a major reduction in Bacilli plus other rarer classes. A detailed investigation of the families belonging to Gammaproteobacteria showed an almost complete dominance of Enterobacteriaceae in the diseased ileum samples, while the control individuals harboured a diverse pattern of different Gammaproteobacteria families (Figure S5).

**Figure 4.**
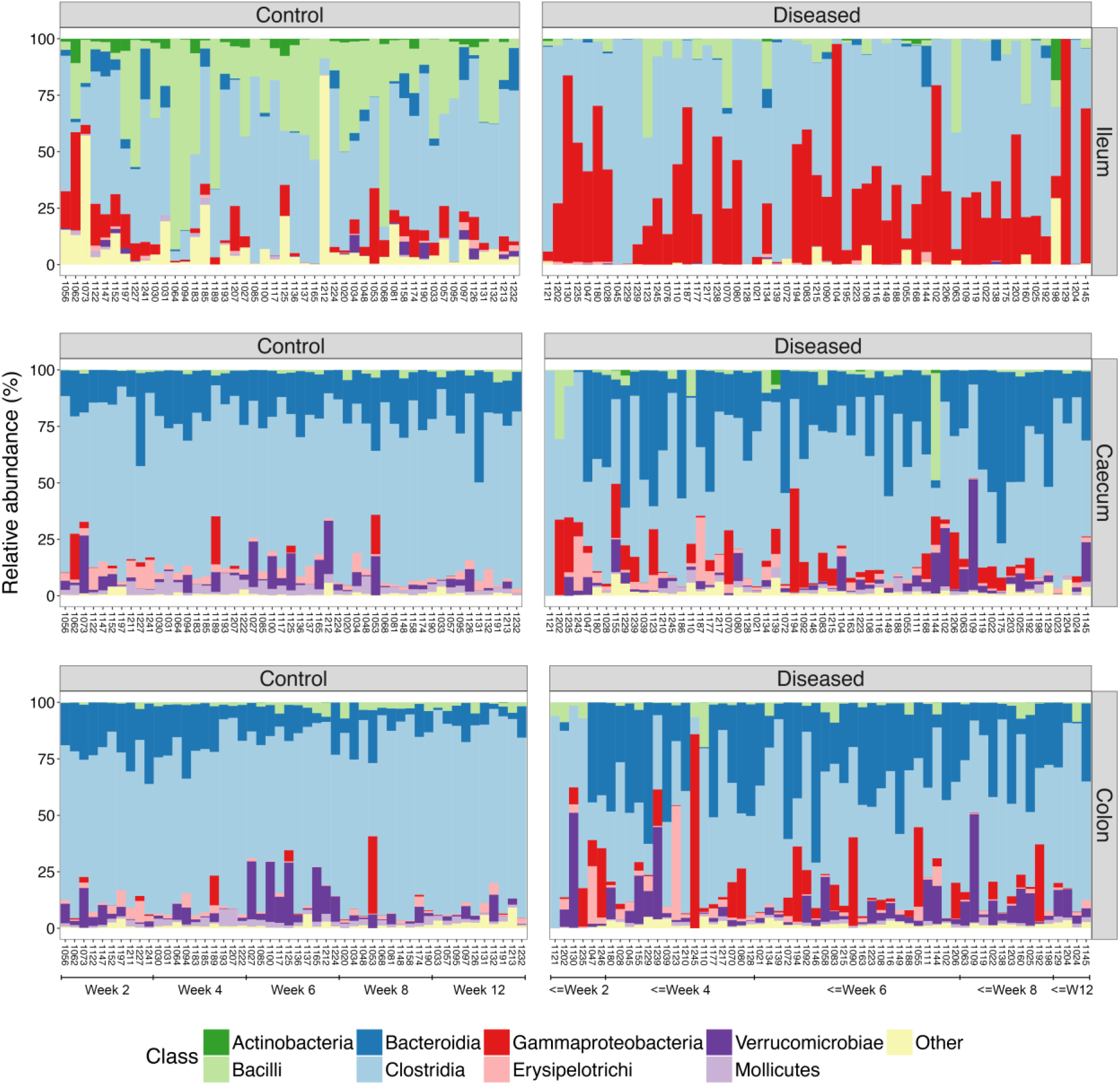
The proportion of bacterial classes per individual and gut region, sorted by age (left bars = youngest, right bars = oldest). Left column = control individuals, right column = diseased individuals. Top row = ileum, middle row = caecum, bottom row = colon.

The Gram-negative Enterobacteriaceae is a large family that is well-known for encompassing several familiar intestinal pathogens and pathobionts, and is frequently seen in higher abundances in hosts with gut dysbiosis (Lupp *et al.* 2007; Garrett *et al.* 2010; Hughes *et al.* 2017). There were 19 Operational Taxonomic Units (OTUs; sequences with 100% nucleotide identity) associated with Enterobacteriaceae in the ileum, and blast searches against the NCBI nucleotide database resulted in matches to a wide range of genera, including *Escherichia*, *Klebsiella*, *Shigella*, *Salmonella*, *Yokenella*, *Citrobacter*, *Enterobacter*, *Cronobacter*, *Antlantibacter*, *Pluralibacter*, *Leclercia*, and *Kluyvera*. In previous studies, it has been shown that various members of the Enterobacteriaceae family often co-occur and bloom simultaneously during dysbiosis (Gevers *et al.* 2014; McDonald *et al.* 2016), which agrees with our findings of multiple genera being present.

Another key characteristic of dysbiosis in the ileum was that certain individuals had microbiomes almost entirely comprised of Clostridia, a pattern not observed in any control individuals (Figure 4). The families of Clostridia showed further striking taxonomic patterns in diseased individuals, including a major increase of Peptostreptococcaceae and a marked reduction of Ruminococcaceae and other rare families (Figure S5). The Peptostreptococcaceae family was represented by six OTUs in our data, and blast searches yielded matches to various species of *Paeniclostridium*, *Paraclostridium*, and *Clostridium*. The most prevalent of these OTUs matched *Paeniclostridium sordellii*, a bacteria known to comprise virulent strains causing high morbidity and mortality through enteritis and enterotoxaemia in both humans and animals (Aldape *et al.* 2006; Sasi Jyothsna *et al.* 2016).

We next identified specific OTUs associated with dysbiosis by performing negative binomial Wald tests of bacterial abundances while controlling for the age of the hosts. Thirty-eight OTUs were significantly overrepresented in the ileum of diseased individuals (Figure 5), of which most belonged to Clostridia, including Ruminococcaceae, various *Clostridium* spp., and *Epulopiscium*, but also *Bacteroides*, *Escherichia*, and *Bilophila wadsworthia* (Table S1).

**Figure 5.**
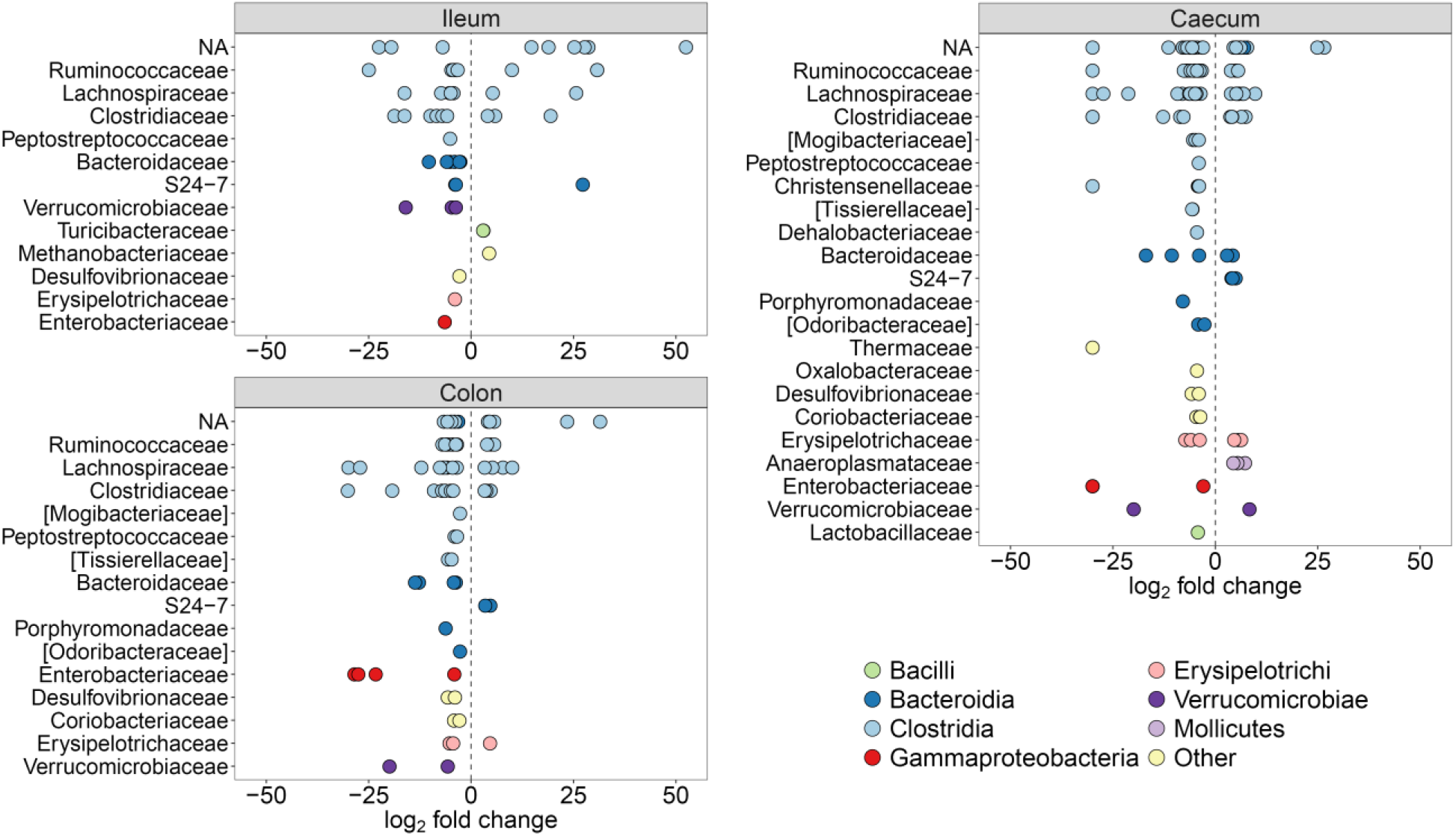
Differentially abundant OTUs (q < 0.01) between control and diseased individuals, separate for the three gut regions. y-axes show taxonomic families and OTUs have been coloured at the class level. Positive log_2_ fold changes indicate higher OTU abundance in the control individuals and negative log_2_ fold changes indicate higher abundance in the diseased individuals. NA = OTUs without family classification.

### Taxa associated with disease in the caecum and colon

The taxonomic composition of the caecum and colon was largely similar across control individuals, exhibiting a relatively stable microbiome composition across hosts and ages. However, both gut regions showed major disruptions in the microbial composition of diseased individuals (Figure 4). Similar to the ileum, the Gammaproteobacteria were again more prevalent in caecal and colon samples of diseased individuals, but a reduction in Clostridia and an increase in Bacteroidia constituted the most prominent differences. Further taxonomic analyses of Bacteroidia showed that the family Porphyromonadaceae had proliferated in diseased individuals (Figure S5). This family encompassed two species in our data, *Parabacteroides distasonis* and *Dysgonomonas* sp. which are found in normal gut microbiota (Sakamoto 2014), However, *P. distasonis* has previously been identified as a colitis-promoting species in mice (Dziarski *et al.* 2016) and *Dysgonomonas* members are known to be associated with cachexia and intestinal inflammation (Huang *et al.* 2017).

Differential abundance tests identified large similarities in the dysbiosis patterns of the caecum and colon, as 50 out of 56 (89%) OTUs that were more abundant in the diseased colon samples were also more abundant in the diseased caecal samples (Figure 5; Tables S2–S3). In addition, 15 out of these OTUs (39%) were also significantly overrepresented in the ileum (Table S1). The most significant OTU in the caecum (q = 1.2e−53) and colon (q = 2.4e−56) was absent in control individuals but abundant in diseased individuals (Tables S2–S3). This OTU, which was also highly significant in the ileum (q = 3.4e−21), had a 100% match against *Clostridium paraputrificum*, a known human pathogen associated with sepsis and necrotizing enterocolitis (Brook & Gluck 1980; Shandera *et al.* 1988; Smith *et al.* 2011). *C. paraputrificum* has also been experimentally studied in gnotobiotic quails (*Coturnix coturnix*), where it caused lesions and haemorrhages in the gut lining associated with enterocolitis (Waligora-Dupriet *et al.* 2005).

Besides *C. paraputrificum*, highly significant OTUs that were more abundant in diseased individuals (Tables S2–S3) gave blast matches (99.5–100% identity) to the *Clostridium* species *C. colinum*, *C. cadaveris*, *C. butyricum*, and *C. perfringens*, all of which have previously been linked to acute inflammation of the gut (enterocolitis) (Ononiwu *et al.* 1978; Snyman *et al.* 1992; Staempfli *et al.* 1992; Shanawany & Dingle 1999; Cassir *et al.* 2016). Other OTUs highly overrepresented in diseased caecal and colon samples belonged to e.g. Enterobacteriaceae, Ruminococcaceae, Mogibacteriaceae, *Bacteroides*, *Dorea*, *Sedimentibacter*, *Bilophila wadsworthia*, and *Eggerthella lenta* (Figure 5; Tables S2–S3). Many of these bacteria constitute part of the normal gut microbiota (Finegold *et al.* 1992; Pan & Yu 2014; David *et al.* 2014), and besides OTUs from *C. paraputrificum*, *C. colinum*, and most Enterobacteriaceae, the majority of all significant OTUs were also present in some control individuals, albeit at much lower abundances (Tables S2–S3).

### Taxa associated with health

The ilea of diseased individuals showed large reductions of certain bacteria compared to controls (Figure 4), mainly Bacilli, a class in which Turicibacteraceae and Lactobacillaceae were the most common families. Turicibacteraceae included two significant OTUs from *Turicibacter* (Table S1), which showed decreased abundances in the diseased ileum. *Turicibacter* has previously been shown to have high heritability in the small intestines of humans and mice where it is in direct contact with host cells, suggesting that it is symbiotic (Goodrich *et al.* 2016). This genus has been associated with both health and disease, but is often found to be depleted in animals with diarrhoea and gut disease (Markel *et al.* 2012; Suchodolski *et al.* 2012; Amato *et al.* 2016).

One of the most striking differences in both the caecum and colon of the diseased individuals was a substantial reduction of the Bacteroidia family S24-7 (Figure S5). Little is known about the S24-7 family, despite being a prominent component of the normal vertebrate gut microbiota (Ormerod *et al.* 2016), but studies of mice have similarly reported a potential beneficial effect of this family, as it is often reduced in diseased hosts (Ferrere *et al.* 2017; Meisel *et al.* 2017). The majority of OTUs with reduced abundance in the colon of diseased individuals were also underrepresented in the caecum (15 out of 19; 79%), indicating large-scale depletion of potentially health-associated bacteria across the hindgut. These OTUs belonged to taxa such as Lachnospiraceae (e.g. *Coprococcus*, *Blautia*), Ruminococcaceae (e.g. *Ruminococcus*), S24-7, Erysipelotrichaceae, *Clostridium*, *Anaeroplasma*, *Turicibacter*, *Methanobrevibacter*, *Akkermansia muciniphila*, and several unknown Clostridiales (Figure 5; Tables S2–S3).

While 15 OTUs were found to be significantly overrepresented in all three gut regions of diseased individuals, only a single OTU was significantly underrepresented in all gut regions of diseased individuals. A blast search of this OTU gave a perfect match to the butyrate-producing genus *Roseburia*, which has repeatedly been associated with health. For example, lower abundances of *Roseburia* spp. have been discovered in humans with ulcerative colitis, inflammatory bowel disease, irritable bowel syndrome, obesity, hepatic encephalopathy, and type 2 diabetes (Bajaj *et al.* 2012; Morgan *et al.* 2012; Machiels *et al.* 2014; Tamanai-Shacoori *et al.* 2017), and in pigs with swine dysentery (Burrough *et al.* 2017). Together, our results suggest that *Roseburia* and many other taxa previously found to be negatively associated with disease, are not specific to mammalian dysbiosis patterns, but are similarly depleted during dysbiosis in phylogenetically distant animals such as ostriches.

### Disruption of the gut microbiota in the weeks preceding death

To establish whether dysbiosis occurs immediately before death or results from imbalances emerging earlier in life, we examined the microbiota of faecal samples repeatedly collected prior to mortality. We found that survival up to four weeks of age was not related to alpha or phylogenetic diversity earlier in life (Table S4). However, mortality after six weeks was related to alpha and phylogenetic diversity when chicks were two to four weeks old. Higher alpha diversity at week two was positively related to survival beyond six weeks of age (Figure S6; Table S4). Conversely, higher alpha diversity at four weeks of age and higher phylogenetic diversity at ages two to four weeks were associated with an increased risk of mortality (Figure S6; Table S4). These results suggest that individuals with low alpha diversity at two weeks of age were susceptible to colonisation of distinct phylogenetic groups of bacteria, which increased the risk of mortality in the subsequent weeks.

Next, we examined if the abundances of bacterial families that differed between diseased and control individuals could predict patterns of future mortality in the weeks leading up to death. There was only weak evidence for any beneficial effects of bacteria with Lactobacillaceae abundance at week two and Turicibacteraceae at week four having a tendency to positively influence survival (Figure S7; Table S4). However, there were very strong associations between the abundance of Peptostreptococcaceae and S24-7 during the first week of life and mortality at all subsequent ages, even after controlling for the abundances of these bacterial families at later ages (Figure 6; Table S4). This result strongly suggests that the time immediately after hatching is a critical period for the establishment of specific pathobionts that can increase the risk of mortality even months later.

**Figure 6.**
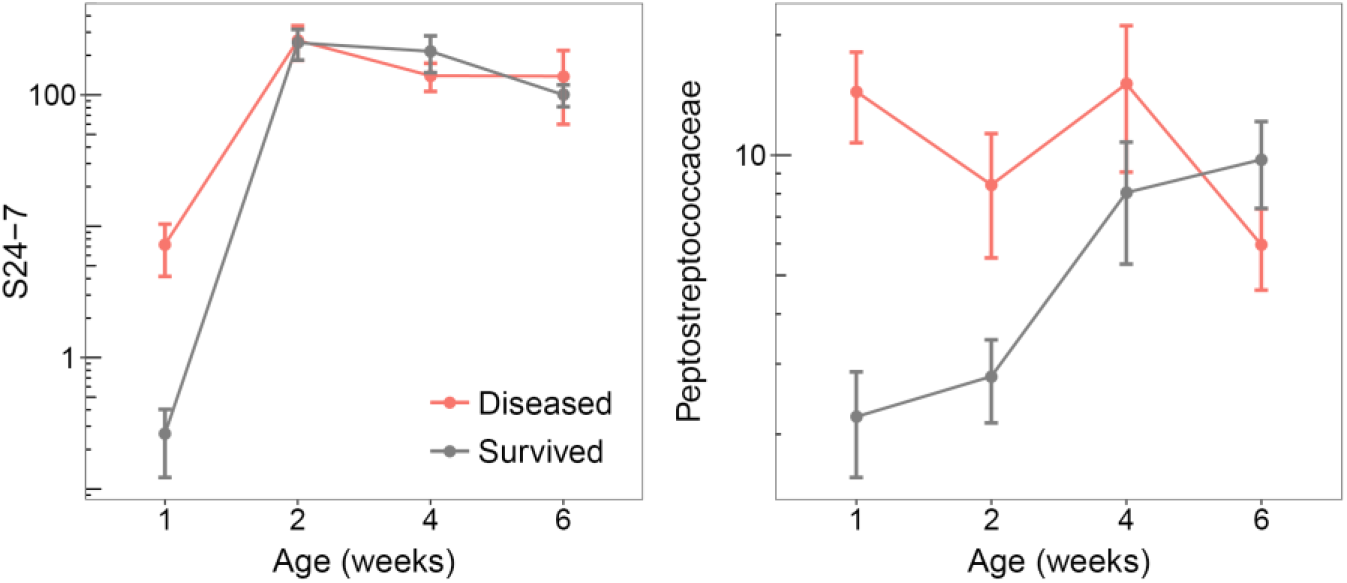
Abundances (normalised and log-transformed) of two bacterial families associated with disease in the weeks preceding death, measured by repeated faecal sampling of individuals. Points and error bars represent means ± SE.

### Environmental sources of gut bacteria

Finally, we evaluated potential environmental sources of microbes present in the gut using SourceTracker. There was essentially no contribution from the water supply (0.1–0.4%) or from the soil (0.2–0.7%) to the gut microbiota of either diseased or control individuals (Figure 7). Instead, the majority of gut bacteria were from unknown sources (89.9%). The microbial sequences of the chicks’ food did overlap with some of the OTUs found in the ileum and colon, but predominantly in control individuals (Figure 7). It is likely that healthy juveniles ate more than sick individuals, and therefore apparently acquired a greater proportion of food-related microbes. These findings indicate that contaminated food or water were not a major source of the bacteria associated with mortality, further supporting the idea that dysbiosis develops from taxa already present in the gut, rather than through acquisition of new taxa.

**Figure 7.**
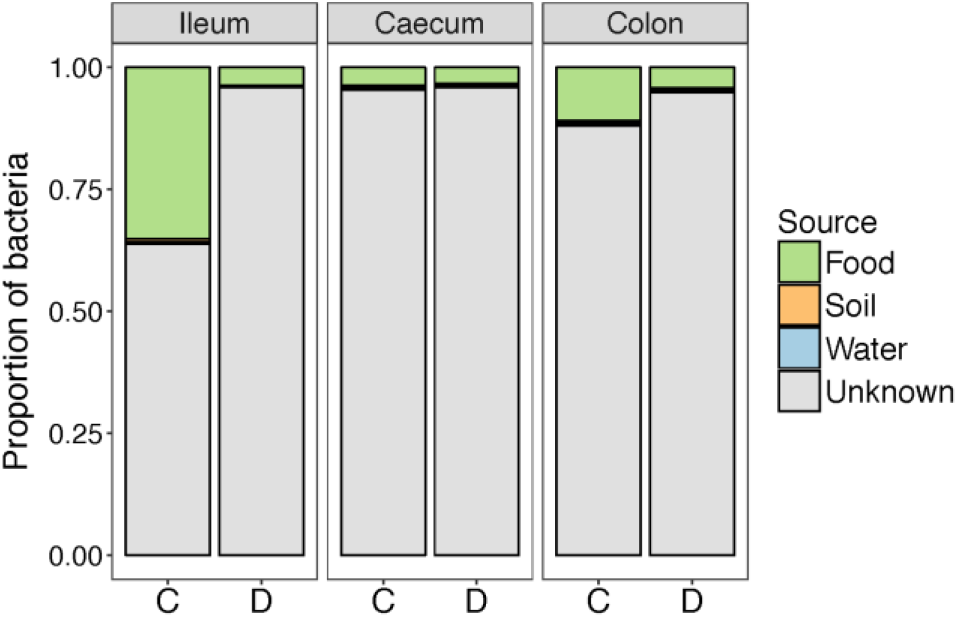
Environmental sources of bacteria present in the different gut sections. C = control individuals and D = diseased individuals.

### Conclusions

Our study shows that severe gut bacterial community disruption is linked to high levels of mortality in developing ostrich chicks. Large-scale shifts in taxon composition, low alpha diversity, and multiple differentially abundant OTUs underlie the dysbiosis pattern seen in diseased individuals. Several taxa associated with disease were disproportionally proliferated in the ileum, caecum, and colon (e.g. Enterobacteriaceae, Peptostreptococcaceae, Porphyromonadaceae, *Clostridium*, *Paeniclostridium*) whereas other taxa were associated with health (e.g. S24-7, Lachnospiraceae including *Roseburia, Coprococcus*, and *Blautia*, Ruminococcaceae, Erysipelotrichaceae, and *Turicibacter*). Dysbiosis was particularly pronounced in the ileum and in individuals that died at early ages, showing that disruptions to gut microbiota develop in a distinct spatial and temporal manner. The establishment of some of the pathogenic bacteria occurred during the initial period after hatching, which predicted patterns of mortality several weeks later. Yet the environment did not show any evidence of pathogenic sources. One striking feature of the dysbiosis we observed is that many of the implicated harmful and beneficial bacteria have been found to have similar effects in a diverse set of vertebrate hosts including humans. This suggests that there is a high degree of evolutionary convergence across some host-microbe associations and that studies on different vertebrate species may contribute to a general understanding of gut dysbiosis.

## Materials and methods

### Experimental setup

We kept ostriches under controlled conditions at the Western Cape Department of Agriculture’s ostrich research facility in Oudtshoorn, South Africa. Eggs were artificially incubated to synchronize hatching on Sep 30^th^ 2014. A total of 234 individuals were reared in four groups of approximately 58 chicks each and monitored from hatching until 12 weeks of age. The groups were kept in indoor pens of approximately 4×8 m in the same building with access to outdoor enclosures with soil substrate during the day, weather permitting. To reduce potential environmental variation on the development of the gut microbiota, all individuals were raised under standardized conditions with *ad libitum* food and water during daytime. The chicks were fed a balanced pelleted ration normally given to ostrich chicks (details in Videvall *et al.* 2019), and were kept in an area completely separate from adult ostriches. No medicine or antibiotics were given to the chicks during this period. All procedures were approved by the Departmental Ethics Committee for Research on Animals (DECRA) of the Western Cape Department of Agriculture, reference no. R13/90.

### Sample collection

A total of 68 individuals died of suspected gut disease during the first three months of age, which we have referred to throughout the text as “diseased”. Multiple chicks exhibited characteristic behaviour of sickness shortly before dying (poor appetite, inactivity, listlessness, depressed posture). All individuals that died were dissected. Additionally, ten chicks (2-3 individuals from each group) were randomly selected for euthanization and dissection at 2, 4, 6, 8, 10, and 12 weeks of age, to act as age-matched controls for the diseased individuals that died. Contents of the ileum, caecum, and colon of all control and diseased individuals were collected in 2 ml micro tubes. To minimize contamination between samples and individuals, lab benches and surfaces were routinely sterilized with 70% ethanol, and equipment used during dissection was cleaned with hot water, 70% ethanol, and placed in the open flame of a Bunsen burner between each sample collection. In addition to intestinal samples, we routinely collected faecal samples at 1, 2, 4, 6, 8, 10, and 12 weeks of age. For more information on the faecal sample collection, please see Videvall *et al.* (2019). Weight measurements of all individuals were obtained at hatching, during each faecal collection event, and immediately before dissection. Environmental samples were collected throughout the experiment by wetting sterile cotton swabs in phosphate-buffered saline (PBS) and swabbing food, drinking water, and the soil of the ostrich chicks’ enclosures. All samples were frozen at −20 °C after collection.

An inflammation score between 0–4 was given each gut region of every individual (n = 323) based on photographs of the gastrointestinal tract of the dissected individuals. The author (E.V.) performing the inflammation assessment was blind to whether individuals had been euthanized or died (control/diseased). The inflammation score was given as follows: 0 = no visible inflammation, 1 = minor inflammation, 2 = intermediate inflammation, 3 = major inflammation, and 4 = extreme and severe inflammation. Twenty-three measures (7%) were given a score of NA because it was not possible to assess the inflammation (e.g. gut region not properly visible) (Table S5).

### DNA sequencing

DNA was extracted using the PowerSoil-htp 96 well soil DNA isolation kit (Mo Bio Laboratories, cat no. 12955-4) as recommended by the Earth Microbiome Project (www.earthmicrobiome.org). Microbial communities were characterized using Illumina MiSeq sequencing to identify the V3 and V4 regions of the 16S rRNA gene. For full details please see Videvall *et al.* (2017, 2018). We sequenced a total of 323 ileum, caecum, and colon samples from all individuals that died (n = 68) and all euthanized (control) individuals at 2, 4, 6, 8, and 12 weeks of age (n = 50 in total; ten individuals per week). We additionally sequenced 460 faecal samples (Videvall *et al.* 2019), 24 environmental samples (food, water, soil), and 4 negative samples (blanks) (Table S5).

### Data processing

Reads were trimmed using Trimmomatic (v. 0.35) (Bolger *et al.* 2014) and quality-filtered in QIIME (v. 1.9.1) (Caporaso *et al.* 2010). Amplicon sequence variants (ASVs, here referred to as operational taxonomic units; OTUs) were clustered in Deblur (v. 1.0.0) (Amir *et al.* 2017) and assigned using the RDP classifier (Wang *et al.* 2007). For further details, please see Videvall *et al.* (2019). We removed all OTUs that were either classified as mitochondria or chloroplast, present in the negative samples, only appeared in one sample, or with a total sequence count of less than 10. These filtering steps together removed approximately 27,000 OTUs, with 5118 remaining for analyses. We further filtered out all samples with a total sequence count of less than 500, resulting in seven ileal and three environmental samples being excluded.

### Data analyses

All statistical analyses were performed in R (v. 3.3.2) (R Core Team 2017) and all plots were made using ggplot2 (Wickham 2009). Bray-Curtis and weighted UniFrac distances between microbiomes were calculated in phyloseq (v. 1.19.1) (McMurdie & Holmes 2013) and examined using a PERMANOVA with the adonis function in vegan (v. 2.4-2) (Oksanen *et al.* 2017). Age effects on the microbiome were evaluated by fitting a linear term and a quadratic age term with Z-transformed values. Beta diversity was tested with a multivariate homogeneity of groups dispersions test using the betadisper function in vegan. We calculated alpha diversity using Shannon’s H index and phylogenetic diversity using Faith’s weighted abundance of phylogenetic diversity. Variation in diversity was analysed using a GLM with a Gaussian error distribution, health status (control versus diseased), age, and their interaction as fixed effects. Separate GLMs were used for each gut region. To evaluate bacterial abundances, we first modelled counts with a local dispersion model and normalised per sample using the geometric mean, according to DESeq2 (Love *et al.* 2014). Differential OTU abundances between control and diseased individuals were subsequently tested in DESeq2 with a negative binomial Wald test while controlling for the age of individuals and with the beta prior set to false (Love *et al.* 2014). Results for the specific comparisons were extracted (e.g. “ileum control” versus “ileum diseased”) and p-values were corrected with the Benjamini and Hochberg false discovery rate for multiple testing. OTUs were labelled significantly differentially abundant if they had an adjusted p-value (q-value) < 0.01. Environmental samples were analysed with SourceTracker (Knights *et al.* 2011).

To estimate the ages at which diversity and bacterial taxa predicted survival we analysed the faecal samples using Cox Proportional Hazards models in survival (v. 2.44-1.1) (Therneau & Grambsch 2000). These models examine whether explanatory variables are associated with a greater risk (beta coefficient >1) or lower risk (beta coefficient < 1) of mortality. Separate models were fitted for each measure of diversity and each bacterial family, for samples collected at 1, 2, 4, and 6 weeks of age and included as explanatory variables. These earlier ages were examined as they had enough mortality data following these time points to estimate effects on survival. Because individuals that died very early in life had missing data for later time points, it was not possible to include all explanatory variables simultaneously without restricting the data to individuals that survived past week 6. Therefore, measures from each age were sequentially entered into models in a chronological order (e.g. week 1 followed by week 1 & 2). By doing this we were able to test how microbiome features at week 1 predicted survival past week 1, how microbiome features at week 2 after controlling for any differences at week 1 predicted survival past week 2, and so forth.

## Data availability

Supporting information has been made available online. Sequences have been uploaded to the European Nucleotide Archive at EBI-EMBL under accession numbers: PRJEB28512 and PRJEB28515.

## Acknowledgements

We are grateful to all staff at the Oudtshoorn Research Farm, Western Cape Government, for assisting with sample collection. Funding was provided by the Helge Ax:son Johnson Foundation, Längmanska Cultural Foundation, Lund Animal Protection Foundation, Lars Hierta Memorial Foundation, and the Royal Physiographic Society of Lund to E.V., by a Wallenberg Academy fellowship (2013.0129) and a Swedish Research Council grant (2017-03880) to C.K.C., and by the Western Cape Government.

## Author contributions

E.V. and C.K.C. planned and designed the study. S.C. provided animal facilities. A.E. supervised the experimental part of the study. N.S., A.E., C.K.C., and E.V. performed the sampling and cared for the animals. A.O. performed the euthanization and advised on sampling. M.S. supervised the laboratorial part of the study, and together with H.M.B. prepared the samples for sequencing. E.V. and C.K.C. performed the bioinformatic and statistical analyses. S.J.S., R.K., and O.H. provided advice on analyses and the interpretation of results. E.V. and C.K.C. wrote the paper with input from all authors.

## References

Aldape MJ, Bryant AE, Stevens DL (2006) Clostridium sordellii Infection: Epidemiology, Clinical Findings, and Current Perspectives on Diagnosis and Treatment. Clinical Infectious Diseases, 43, 1436–1446.

Amato KR, Metcalf JL, Song SJ et al. (2016) Using the gut microbiota as a novel tool for examining colobine primate GI health. Global Ecology and Conservation, 7, 225–237.

Amir A, McDonald D, Navas-Molina JA et al. (2017) Deblur Rapidly Resolves Single-Nucleotide Community Sequence Patterns. mSystems, 2, e00191–16.

Antharam VC, Li EC, Ishmael A et al. (2013) Intestinal dysbiosis and depletion of butyrogenic bacteria in *Clostridium difficile* infection and nosocomial diarrhea. Journal of Clinical Microbiology, 51, 2884–2892.

Bajaj JS, Hylemon PB, Ridlon JM et al. (2012) Colonic mucosal microbiome differs from stool microbiome in cirrhosis and hepatic encephalopathy and is linked to cognition and inflammation. AJP: Gastrointestinal and Liver Physiology, 303, G675–G685.

Berer K, Gerdes LA, Cekanaviciute E et al. (2017) Gut microbiota from multiple sclerosis patients enables spontaneous autoimmune encephalomyelitis in mice. Proceedings of the National Academy of Sciences, 114, 10719–10724.

Bolger AM, Lohse M, Usadel B (2014) Trimmomatic: A flexible trimmer for Illumina sequence data. Bioinformatics, 30, 2114–2120.

Brook I, Gluck RS (1980) *Clostridium paraputrificum* sepsis in sickle cell anemia. Southern Medical Journal, 73, 1644–1645.

Burrough ER, Arruda BL, Plummer PJ (2017) Comparison of the Luminal and Mucosa-Associated Microbiota in the Colon of Pigs with and without Swine Dysentery. Frontiers in Veterinary Science, 4.

Caporaso JG, Kuczynski J, Stombaugh J et al. (2010) QIIME allows analysis of high-throughput community sequencing data. Nature Methods, 7, 335–336.

Cassir N, Benamar S, La Scola B (2016) *Clostridium butyricum*: From beneficial to a new emerging pathogen. Clinical Microbiology and Infection, 22, 37–45.

Chow J, Tang H, Mazmanian SK (2011) Pathobionts of the gastrointestinal microbiota and inflammatory disease. Current Opinion in Immunology, 23, 473–480.

Cloete SWP, Brand TS, Hoffman L et al. (2012) The development of ratite production through continued research. World’s Poultry Science Journal, 68, 323–334.

Cloete SWP, Lambrechts H, Punt K, Brand Z (2001) Factors related to high levels of ostrich chick mortality from hatching to 90 days of age in an intensive rearing system. Journal of the South African Veterinary Association, 72, 197–202.

David LA, Maurice CF, Carmody RN et al. (2014) Diet rapidly and reproducibly alters the human gut microbiome. Nature, 505, 559–563.

Donaldson GP, Lee SM, Mazmanian SK (2015) Gut biogeography of the bacterial microbiota. Nature Reviews Microbiology, 14, 20–32.

Duvallet C, Gibbons SM, Gurry T, Irizarry RA, Alm EJ (2017) Meta-analysis of gut microbiome studies identifies disease-specific and shared responses. Nature Communications, 8, 1784.

Dziarski R, Park SY, Kashyap DR, Dowd SE, Gupta D (2016) Pglyrp-Regulated Gut Microflora *Prevotella falsenii*, *Parabacteroides distasonis* and *Bacteroides eggerthii* Enhance and *Alistipes finegoldii* Attenuates Colitis in Mice. PLoS ONE, 11, e0146162.

Ferrere G, Wrzosek L, Cailleux F et al. (2017) Fecal microbiota manipulation prevents dysbiosis and alcohol-induced liver injury in mice. Journal of Hepatology, 66, 806–815.

Finegold S, Summanen P, Hunt Gerardo S, Baron E (1992) Clinical importance of *Bilophila wadsworthia*. European Journal of Clinical Microbiology and Infectious Diseases, 11, 1058–1063.

Frazier KS, Herron AJ, Hines ME, Gaskin JM, Altman NH (1993) Diagnosis of enteritis and enterotoxemia due to *Clostridium difficile* in captive ostriches (*Struthio camelus*). Journal of Veterinary Diagnostic Investigation, 5, 623–625.

Garrett WS, Gallini CA, Yatsunenko T et al. (2010) Enterobacteriaceae act in concert with the gut microbiota to induce spontaneous and maternally transmitted colitis. Cell Host and Microbe, 8, 292–300.

Gevers D, Kugathasan S, Denson LA et al. (2014) The Treatment-Naive Microbiome in New-Onset Crohn’s Disease. Cell Host & Microbe, 15, 382–392.

Goodrich JK, Davenport ER, Waters JL, Clark AG, Ley RE (2016) Cross-species comparisons of host genetic associations with the microbiome. Science, 352, 532–535.

Huang G, Khan I, Li X et al. (2017) Ginsenosides Rb3 and Rd reduce polyps formation while reinstate the dysbiotic gut microbiota and the intestinal microenvironment in ApcMin/+ mice. Scientific Reports, 7, 12552.

Huchzermeyer FW (1999) Veterinary problems. In: The ostrich: biology, production and health (ed Deeming DC), pp. 293–320.

Hughes ER, Winter MG, Duerkop BA et al. (2017) Microbial Respiration and Formate Oxidation as Metabolic Signatures of Inflammation-Associated Dysbiosis. Cell Host & Microbe, 21, 208–219.

Keokilwe L, Olivier A, Burger WP et al. (2015) Bacterial enteritis in ostrich (*Struthio Camelus*) chicks in the Western Cape Province, South Africa. Poultry Science, 94, 1177–1183.

Klingensmith NJ, Coopersmith CM (2016) The Gut as the Motor of Multiple Organ Dysfunction in Critical Illness. Critical Care Clinics, 32, 203–212.

Knights D, Kuczynski J, Charlson ES et al. (2011) Bayesian community-wide culture-independent microbial source tracking. Nature Methods, 8, 761–763.

Latorre M, Krishnareddy S, Freedberg DE (2015) Microbiome as mediator: Do systemic infections start in the gut? World Journal of Gastroenterology, 21, 10487–10492.

Love MI, Huber W, Anders S (2014) Moderated estimation of fold change and dispersion for RNA-seq data with DESeq2. Genome Biology, 15, 550.

Lupp C, Robertson ML, Wickham ME et al. (2007) Host-Mediated Inflammation Disrupts the Intestinal Microbiota and Promotes the Overgrowth of Enterobacteriaceae. Cell Host and Microbe, 2, 119–129.

Machiels K, Joossens M, Sabino J et al. (2014) A decrease of the butyrate-producing species *Roseburia hominis* and *Faecalibacterium prausnitzii* defines dysbiosis in patients with ulcerative colitis. Gut, 63, 1275–1283.

Markel ME, Berghoff N, Unterer S et al. (2012) Characterization Of Fecal Dysbiosis In Dogs With Chronic Enteropathies And Acute Hemorrhagic Diarrhea. Journal of Veterinary Internal Medicine, 26, 765–766.

McDonald D, Ackermann G, Khailova L et al. (2016) Extreme Dysbiosis of the Microbiome in Critical Illness. mSphere, 1, e00199–16.

McFall-Ngai M, Hadfield MG, Bosch TCG et al. (2013) Animals in a bacterial world, a new imperative for the life sciences. Proceedings of the National Academy of Sciences, 110, 3229–3236.

McMurdie PJ, Holmes S (2013) Phyloseq: An R Package for Reproducible Interactive Analysis and Graphics of Microbiome Census Data. PLoS ONE, 8, e61217.

Meisel M, Mayassi T, Fehlner-Peach H et al. (2017) Interleukin-15 promotes intestinal dysbiosis with butyrate deficiency associated with increased susceptibility to colitis. ISME Journal, 11, 15–30.

Morgan XC, Tickle TL, Sokol H et al. (2012) Dysfunction of the intestinal microbiome in inflammatory bowel disease and treatment. Genome Biology, 13, R79.

Oksanen J, Blanchet FG, Friendly M et al. (2017) vegan: Community Ecology Package. R package version 2.4-2.

Ononiwu JC, Prescott JF, Carlson HC, Julian RJ (1978) Ulcerative enteritis caused by *Clostridium colinum* in chickens. The Canadian Veterinary Journal, 19, 226–229.

Ormerod KL, Wood DLA, Lachner N et al. (2016) Genomic characterization of the uncultured Bacteroidales family S24-7 inhabiting the guts of homeothermic animals. Microbiome, 4, 36.

Pan D, Yu Z (2014) Intestinal microbiome of poultry and its interaction with host and diet. Gut Microbes, 5, 108–119.

Petersen C, Round JL (2014) Defining dysbiosis and its influence on host immunity and disease. Cellular Microbiology, 16, 1024–1033.

R Core Team (2017) R: A language and environment for statistical computing. R Foundation for Statistical Computing, Vienna, Austria.

Sakamoto M (2014) The Family Porphyromonadaceae. In: The Prokaryotes (eds Rosenberg E, DeLong EF, Lory S, Stackebrandt E, Thompson F), pp. 811–824. Springer, Berlin, Heidelberg.

Sasi Jyothsna TS, Tushar L, Sasikala C, Ramana CV (2016) *Paraclostridium benzoelyticum* gen. nov., sp. nov., isolated from marine sediment and reclassification of *Clostridium bifermentans* as *Paraclostridium bifermentans* comb. nov. Proposal of a new genus *Paeniclostridium* gen. nov. to accommodate *Clostridium sordellii* and *Clostridium ghonii*. International Journal of Systematic and Evolutionary Microbiology, 66, 1268–1274.

Sekirov I, Russell SL, Antunes LCM, Finlay BB (2010) Gut Microbiota in Health and Disease. Physiological Reviews, 90, 859–904.

Shanawany M., Dingle J (1999) Ostrich production systems (Vol. 144). Food & Agriculture Org.

Shandera WX, Humphrey RL, Stratton LB (1988) Necrotizing enterocolitis associated with *Clostridium paraputrificum* septicemia. Southern Medical Journal, 81, 283–284.

Smith B, Bodé S, Petersen BL et al. (2011) Community analysis of bacteria colonizing intestinal tissue of neonates with necrotizing enterocolitis. BMC Microbiology, 11.

Snyman AE, De Wet SC, Kellerman GE (1992) Clostridial enterotoxemia in young ostriches. In: Proceedings of the South African Veterinary Association Biennial National Congress, p. 185. South African Veterinary Association, Pretoria, Grahamstown, South Africa.

Sommer F, Anderson JM, Bharti R, Raes J, Rosenstiel P (2017) The resilience of the intestinal microbiota influences health and disease. Nature Reviews Microbiology, 15, 630–638.

Staempfli HR, Prescott JF, Carman RJ, McCutcheon LJ (1992) Use of bacitracin in the prevention and treatment of experimentally-induced idiopathic colitis in horses. Canadian Journal of Veterinary Research, 56, 233.

Suchodolski JS, Markel ME, Garcia-Mazcorro JF et al. (2012) The Fecal Microbiome in Dogs with Acute Diarrhea and Idiopathic Inflammatory Bowel Disease. PLoS ONE, 7.

Tamanai-Shacoori Z, Smida I, Bousarghin L et al. (2017) *Roseburia* spp.: a marker of health? Future Microbiology, 12, 157–170.

Therneau TM, Grambsch PM (2000) Modeling Survival Data: Extending the Cox Model. Springer, New York.

Verwoerd DJ (2000) Ostrich diseases. Revue scientifique et technique (International Office of Epizootics), 19, 638–661.

Verwoerd DJ, Deeming DC, Angel CR, Perelman B (1999) Rearing environments around the world. In: The ostrich: biology, production and health (ed Deeming DC), pp. 163–206.

Videvall E, Song SJ, Bensch HM et al. (2019) Major shifts in gut microbiota during development and its relationship to growth in ostriches. Molecular Ecology, 28, 2653–2667.

Videvall E, Strandh M, Engelbrecht A, Cloete S, Cornwallis CK (2017) Direct PCR Offers a Fast and Reliable Alternative to Conventional DNA Isolation Methods for Gut Microbiomes. mSystems, 2, e00132–17.

Videvall E, Strandh M, Engelbrecht A, Cloete S, Cornwallis CK (2018) Measuring the gut microbiome in birds: Comparison of faecal and cloacal sampling. Molecular Ecology Resources, 18, 424–434.

Vijay-Kumar M, Aitken JD, Carvalho FA et al. (2010) Metabolic Syndrome and Altered Gut Microbiota in Mice Lacking Toll-Like Receptor 5. Science, 328, 228–231.

Waligora-Dupriet A-J, Dugay A, Auzeil N, Huerre M, Butel M-J (2005) Evidence for Clostridial Implication in Necrotizing Enterocolitis through Bacterial Fermentation in a Gnotobiotic Quail Model. Pediatric Research, 58, 629–635.

Wang Q, Garrity GM, Tiedje JM, Cole JR (2007) Naïve Bayesian classifier for rapid assignment of rRNA sequences into the new bacterial taxonomy. Applied and Environmental Microbiology, 73, 5261–5267.

Wickham H (2009) ggplot2: elegant graphics for data analysis. New York: Springer.

Wu H-J, Wu E (2012) The role of gut microbiota in immune homeostasis and autoimmunity. Gut Microbes, 3, 4–14.

Zhang Z, Geng J, Tang X et al. (2014) Spatial heterogeneity and co-occurrence patterns of human mucosal-associated intestinal microbiota. The ISME Journal, 8, 881–893.

Zimmermann K, Haas A, Oxenius A (2012) Systemic antibody responses to gut microbes in health and disease. Gut Microbes, 3, 42–47.

